# An annual gene expression profile of markers for programmed cell death and polyphenols biosynthesis across heartwood forming stems in *Robinia pseudoacacia*

**DOI:** 10.1101/2023.10.12.560861

**Authors:** Sylas Lau, Henrik Lange, Elisabeth Magel

**Affiliations:** Wood Biology, Department of Wood Science, University of Hamburg, Leuschnerstraße 91d, 21031 Hamburg, Germany

**Keywords:** Heartwood formation, stem development, programmed cell death, vacuolar processing enzyme (VPE), Metacaspase (MC)

## Abstract

Heartwood formation is a developmental defense strategy in trees that transforms the inner stem core through the deposition of natural antimicrobial preservatives. The defining cellular events in this process are the accumulation of specialized metabolites conjoined with the death of living cells in the transition zone. While previous studies have characterized heartwood compounds and deposition, the cell death aspects are often neglected. Thus, the correlation of these defining cellular events remained unclear, while the exact timing of occurrence and regional involvement within the stem became topics for debate. In this study, members of metacaspases and vacuolar processing enzymes were isolated as programmed cell death (PCD) markers to explore their correlation with those of polyphenol biosynthesis. We collected heartwood-forming stems from *R. pseudoacacia* monthly for a year and monitored their expression through real-time PCR. Our results revealed novel markers of PCD in xylem parenchyma cells and demonstrated their corresponding fluctuation in expression with markers of polyphenols across the stem regions. The annual expression profile demonstrated regional stem development along with weather data on a monthly scale. This work provides a framework to compare the xylem-specific PCD in stages of stem development and a preliminary overview of how they correspond to external conditions over an annual cycle.

## Introductions

The final stage of stem development in many trees is the fortification of the inner stem core with antimicrobial specialized metabolites, a process known as heartwood formation (HWF) ^1,2^. This strengthens the trees’ internal structure and protects them against decay-causing organisms, thereby significantly contributing to their resilience and longevity ^3–5^. Regarding economic benefits, heartwood has superior physical properties as timers and the wide arrays of specialized chemicals from their extractives offer significant values in multiple industries ^6^. HWF is carried out by the xylem ray parenchyma (XP) cells at the transition zone (TZ), and the defining cellular changes are the biosynthesis of heartwood substances (HWS) and the cessation of living cells ^2^. While this development is generally understood to occur in fall and winter ^1,7^, the exact timing has been ambiguous and its molecular regulation has remained enigmatic. Recent studies pointed out the various cellular events during the complex differentiation process in separate seasons ^8,9^. Furthermore, a new theory with genomic evidence challenged the definitions of TZ and HW at the cellular level ^10^, which have been the basis for identifying relevant specimens in biological experiments. Hence, this study closely examines the defining cellular changes of HWF from the perspective of terminal differentiation observed during the woody stem development.

Xylem cell differentiation occurs at the vascular cambium with vigorous production of secondary cell wall (SCW) components and programmed cell death (PCD) ^11,12^. In angiosperms, vessel elements develop into hollow pipe-shaped conduits and fibers mature into brick-like supportive structures ^13,14^. Both cell types are dead at maturity; their remnants, composed of SCW with lignin being the chief constituent, establish the infrastructure for long-distance water conduction ^15^. XP cells remain alive after initial differentiation and are spatially arranged as an interwoven network in the stem, performing essential functions in response to environmental clues ^16–18^. This includes regulating the sap content in the conduits ^19–21^, managing carbon resources gathered from photosynthesis ^22^, and providing dynamic responses to damages and infections ^23–25^. In heartwood-forming trees, after the peak of the vegetative season, carbon reserves in the form of fats and starch stored in the XP cells at the outer stem are broken down into simple sugars and transported to the inner stem. This fuels the production of antimicrobial metabolites by the aged XP cells at the TZ before their imminent death ^26,27^, imparting passive defense to the dying region that is losing active protection ^2,28,29^.

Lignin and HWS share part of their biosynthetic pathway and the activities of enzymes on the diverging route can serve as markers of polyphenol productivity. Wood formation involves the regulation of thousands of genes, which is orchestrated with meticulous transcriptional control ^30^. Transcriptional initiation of relevant biosynthetic enzymes and PCD are directed by the interactions of transcription factors (TFs) such as wood-associated MYB master switches ^31,32^. HWS compose diverse arrays of secondary metabolites and commonly include a prominent profile of flavonoids ^1,2^, which are produced via the general phenylpropanoid biosynthetic pathway that precedes the splitting routes for the production of lignin monomers and flavonoids ^27^ (Figure 1a). Phenylalanine ammonia-lyase (PAL; EC 4.3.1.5) is the initial enzyme in the phenylpropanoid pathway, and chalcone synthase (CHS; EC 2.3.1.74) makes the first committed enzyme in flavonoid biosynthesis ^33^; they are regulated by a comparable set of TFs ^34–37^ (Figure 1b). Evidence found in *R. pseudoacacia* showed a direct correlation between transcription levels of *RpCHS* with flavonols accumulation in the TZ, and thus a key indicator of HWS production ^4^. Moreover, gene expression studies in the same species demonstrated that the isoforms *RpPAL1* and *RpCHS3* were the most prominent markers for polyphenol biosynthesis during HWF ^38^. Thus, these two are employed as references in the current experiment to explore their correlation with PCD markers.

**Figure 1:**
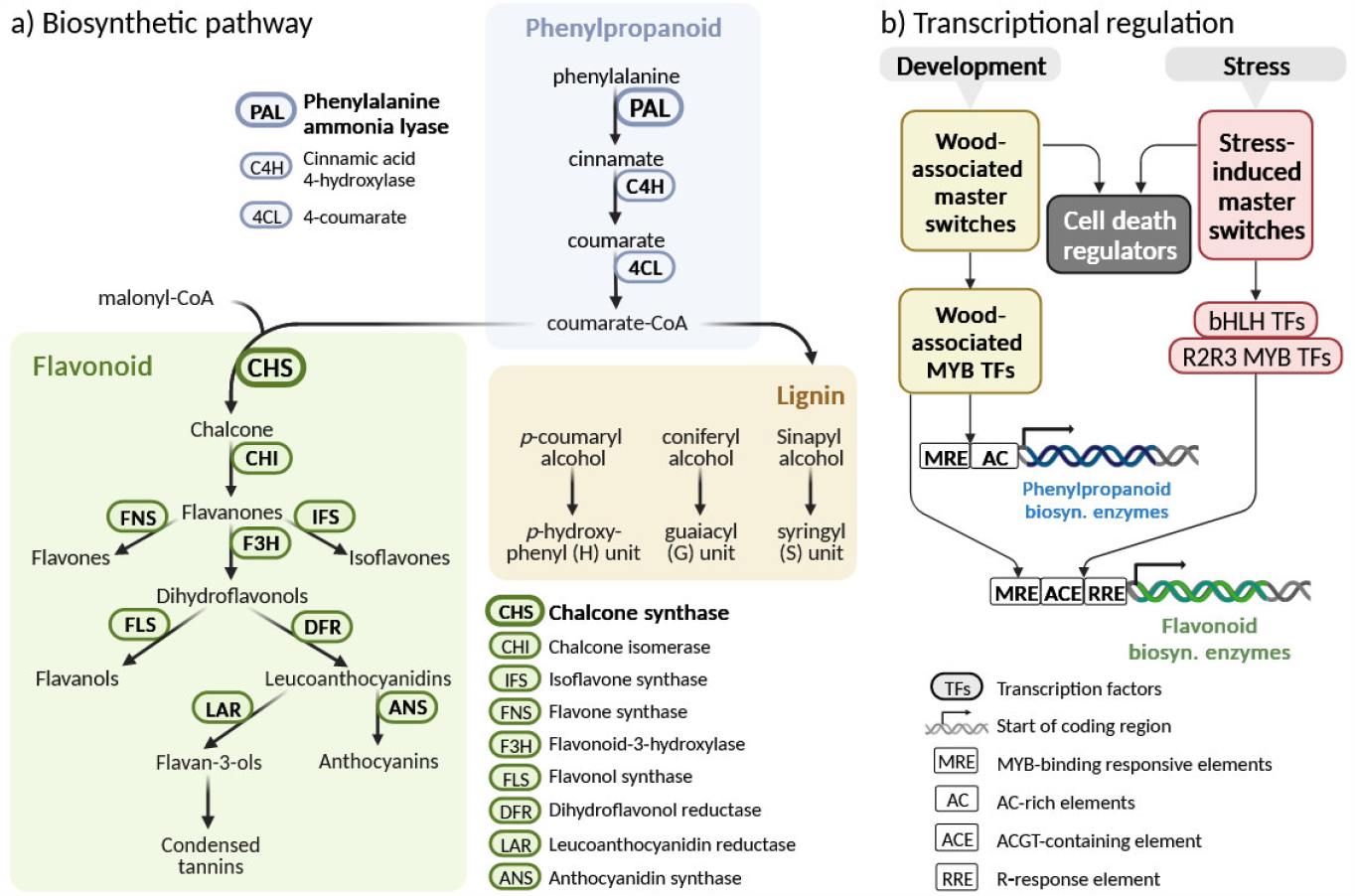
Polyphenol biosynthesis in woody plants. a) Metabolic pathways: Lignin and flavonoid biosynthesis in woody plants share initial steps and precursor molecules within the phenylpropanoid pathway. b) Transcriptional regulation: Specific cis-regulatory elements in the genes of phenylpropanoid and flavonoid biosynthetic enzymes are regulated through combinations of transcription factors (TFs), that are responsive to both developmental and stress-related cues. Wood-associated NAC and MYB master switches are upstream TFs that coordinate the expression of biosynthetic enzymes and programmed cell death. Upstream TFs, so-called master switches, are responsive to developmental and stress-related cues, and they coordinate the expression of biosynthetic enzymes and programmed cell death.

**Figure 2:**
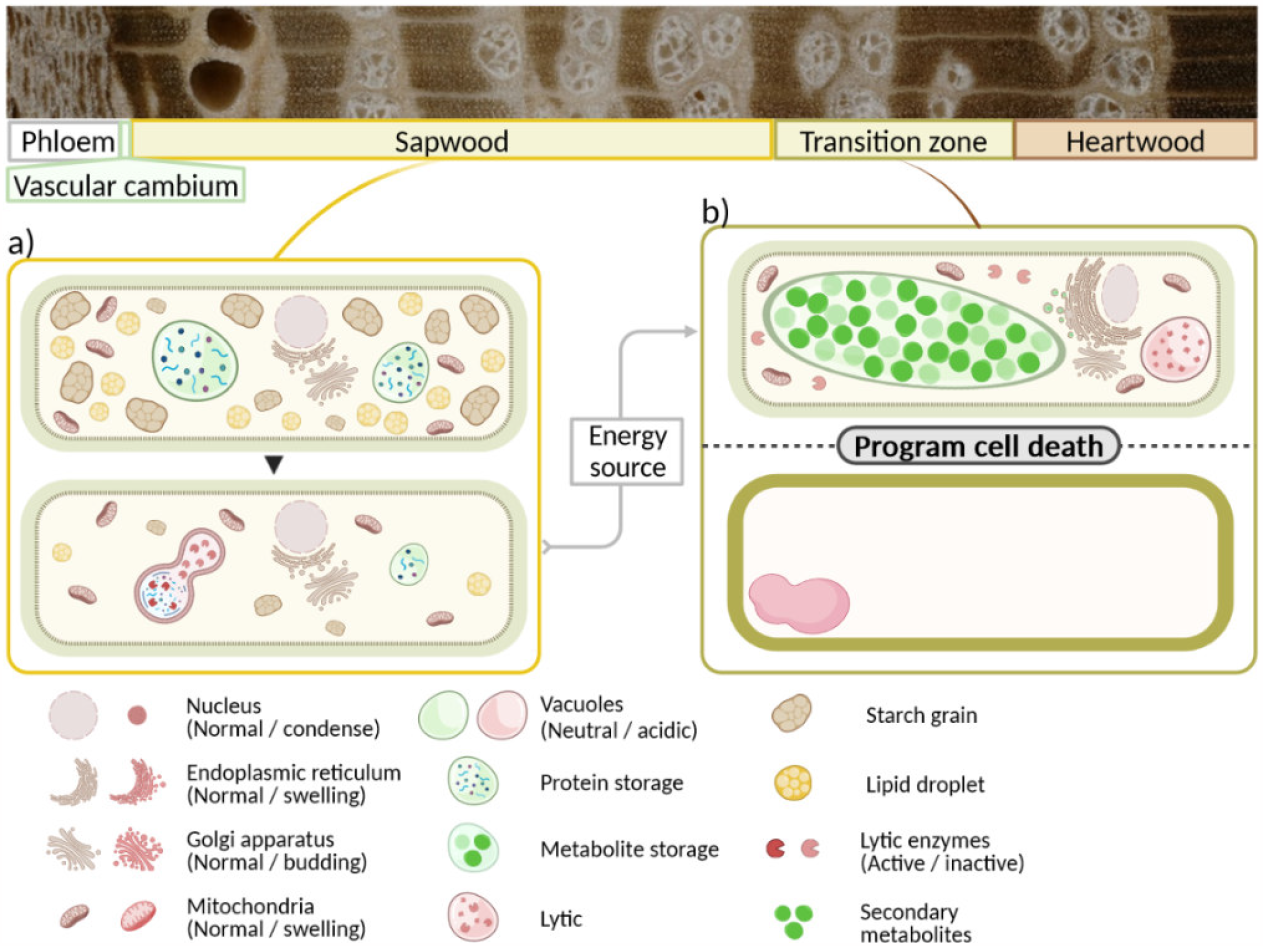
Schematic illustration of the cellular changes in xylem ray parenchyma (XRP) cells during heartwood formation. a) XRP cells in the sapwood. As heartwood formation begins, reserve materials are broken down into simple sugar and transported to the transition zone (TZ). b) XRP cells at the TZ. Vigorous production of phenolic metabolites and zymogens of lytic enzymes are compartmentalized in respective vacuoles. Programmed cell death occurs, the cytoplasm along with organelles are degraded, and phenolic metabolites are deposited into the cell wall matrix. (Image created with BioRender.com)

PCD is highly conserved among multicellular organisms and primarily executed through proteolytic cascades, while autolysis is a central feature observed in plants that marks the moment of death cells. In animal cells, the caspase family interacts with hundreds of substrates to regulate the proteolysis process during PCD ^39^, and metacaspases (MCs; EC 4.3.22.-) are their structural homologs in plants. However, they differ in upstream factors, substrate specificity, and roles in the proteolytic cascade ^40^. Activation of MCs are regulated by specific cellular conditions ^40,41^, such as calcium ions (Ca^2+^) levels and pH optima ^41–43^. As demonstrated in *A. thaliana*, an influx of Ca^2+^ into damaged cells activates MC4, and subsequently initiates the downstream elements for immune response ^44^. Autolysis, ubiquitously observed in plants and considered to mark the moment of cell death, is the collapse of the central vacuole that leads to cytoplasmic degradation ^45,46^. Lytic vacuoles are acidic and contain hydrolytic enzymes; their collapse releases the hydrolases into the cytoplasm and renders a drop in pH, which provides the activating condition for cytosolic enzymes for *post-mortem* degradation. Microarray analysis revealed the upregulation of MC9 during the late stage of TE differentiation in *Arabidopsis* ^47^, while two homologs in hybrid aspen were shown with differential expression during xylem maturation ^48^.

Asparaginyl endopeptidases (AEPs) exhibit caspases-like activities and were identified as the regulators for initiating autolytic cell death in plants ^49,50^. AEPs in mammals are termed legumain and mainly function in the lysosomes; in plants, they are named vacuole processing enzymes (VPEs) for their predominant role in protein processing within the vacuoles ^51,52^. They cleave their substrates after aspartic acid in the YVAD sequence like caspase-1 ^49,50^, in addition to asparagine. Zymogens of VPE encode a signal peptide for them to be transported to specific sub-cellular locations, often the vacuoles, and a C-terminal peptide for auto-activation in an acidic environment ^53,54^. This enables them to act as priming hydrolases in the vacuoles, leading to the proteolytic cascades that cause tonoplast rupture and subsequent degradation of cytoplasmic structures ^53,55^. Upregulation of *VPEs* have been detected during TE differentiation in *Arabidopsis* ^47^ and *Populus* ^56^. Recent research that explored the role of γVPE in stem development revealed its maturation of a papain-like cysteine proteinase CEP1 makes a direct link to secondary walls thickening in xylem fiber cells ^57^. Furthermore, among the list of differentially expressed genes (DEGs) in the *γvpe* mutant during xylogenesis are key enzymes in the lignin monomer synthesis pathway, peroxidases and laccases responsible for monolignols polymerization ^57^.

The present study aims to identify molecular markers defining cellular events during HWF and examine their correlation through gene expression analysis. Based on current knowledge of the cellular process in xylogenesis and HWF, the hypothesis is that the molecular linkage between polyphenols biosynthesis and PCD is a shared characteristic during the terminal differentiation among different xylem cell types. MCs and VPEs, known to be involved in xylem-specific developmental PCD, are selected as markers to facilitate determining the correlation with polyphenol biosynthesis and drawing comparisons between different major stages of stem development.

## Materials and Methods

### Plant materials

Six *R. pseudoacacia* trees grown in the arboretum of the Thünen Institute in Hamburg-Bergedorf, Germany were selected for a full-year sampling regime. They were about 25 years old and 15 to 20 meters in height. In the first week of every month between April 2019 to March 2020, a healthy branch with a 5 to 20 cm diameter was harvested, immediately flash-frozen with dry ice, and transferred to a -80°C freezer. Fully frozen logs were sawn into 2 cm thick disks and lyophilized, then sectioned under a magnifying glass into regions as indicated in Table 1 and Figure 3, carefully avoiding wound wood. For the homogenized sample, 20g of shavings were collected from each region and milled into flour using a Retsch Mill while kept cold with liquid nitrogen.

**Table 1:**
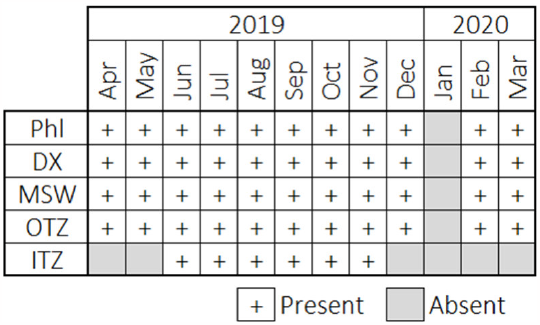
Availability of sample for the present study.

**Figure 3:**
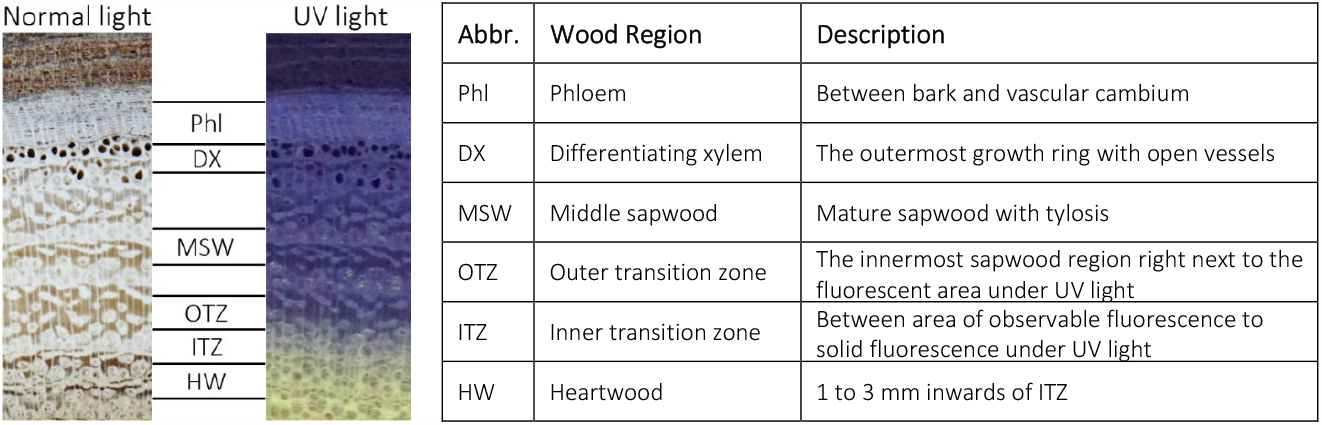
Wood regions for sample sectioning. Photos of a freeze-dried specimen under normal and UV light with indication on the different regions sectioned for the present study. RNA derived from the heartwood region was not sufficient for the downstream process and thus omitted.

### Isolation of candidate genes in *R. pseudoacacia*

Genomic DNA of the samples was extracted using Qiagen DNeasy Plant Kit (Qiagen, Hilden, Germany). Homologous gene sequences from Fabales species were retrieved from NCBI and UniProt databases, and aligned in Geneious Prime ^58^. Degenerate primers were designed based on the sequence alignment in the same software, following guideline recommendations ^59^. PCR was conducted on Biometra thermocyclers T-Gradient (Biomedizinische Analytik GmbH, Göttingen, Germany), using the QIAGEN Taq PCR Core Kit (Qiagen, Hilden, Germany) and primers manufactured by biomers.net GmbH. PCR products were purified using the QIAGEN PCR Purification kit and sent to Eurofins MWG Operon for Sanger sequencing.

### Bioinformatics

Results from Sanger sequencing were imported into Geneious Prime ^58^ for identification with NCBI, assembly, alignment, and primer design. The same software was used for translation into putative protein sequence, and prediction of functional areas with built-in InterPro: Integrative Signature database. 3D structural homology models were generated using the SWISS-MODEL workspace ^60^ based on sequence alignments with homologs and visualized using PyMOL Molecular Graphic System ^61^ (templates accessions and SWISS-MODEL evaluation scores of each model are provided in the supplementary materials).

### Extraction of total RNA and cDNA synthesis

Total RNA was extracted using the RNeasy Plus Universal Kits (Qiagen, Hilden, Germany), following the manufacturer’s instructions. Removal of genomic DNA and cDNA synthesis was carried out using the QuantiNova Reverse Transcription Kit (Qiagen, Hilden, Germany) with 200 ng of total RNA as a template. The cDNA obtained was made into aliquots for qPCR and stored at -80°C until used and -20°C for temporary storage during the experiment.

### Gene expression analysis

qPCR was conducted using the thermal cycler Stratagene MX3005P (Agilent Technologies), assays were performed with triplicates of cDNA aliquots in 96-wells reaction plates using Quantinova qPCR SYBR Green Master Mix (QIAGEN) according to the manufacturer’s manual. Primers were manufactured by biomers.net GmbH, details are listed in supplementary materials. Melting curve analysis was included in each run, and reactions that did not fulfill the primers’ specificity were omitted before the Ct values were averaged. *Rp18S* was previously validated as the reference gene with stable expression during HWF ^38^ and was used for normalization of the threshold cycles (Cq). The analysis of relative expression was carried out with the delta-delta Ct method ^62^.

## Results

### Sequence analysis and structure prediction of the candidate gene in R. pseudoacacia

Four partial gene sequences of type-II MCs (MC-II) were isolated from *R. pseudoacacia*, designated as *RpMC9, RpMC4a, RpMC4b*, and *RpMC4c*. The former is a homolog of *AtMC9*, while the others share high homology with *MC4-like* genes. Except for *RpMC4c*, their translated sequences span over the key residues for comparison with homologs, and contain the conserved aspartates predicted to contribute to substrate specificity (Figure 4). RpMC4a and -b carry a region of negatively charged residues for Ca^2+^-dependent activation. While the AlphaFold DB models of homologs in *Glycine max* share the highest sequence identity in alignment with RpMC-II, protein structure predictions were generated based on the homologs in *A. thaliana* with crystal structures as templates. They showed high homology in the general structures with superimposable helixes, and differ in the coils.

**Figure 4:**
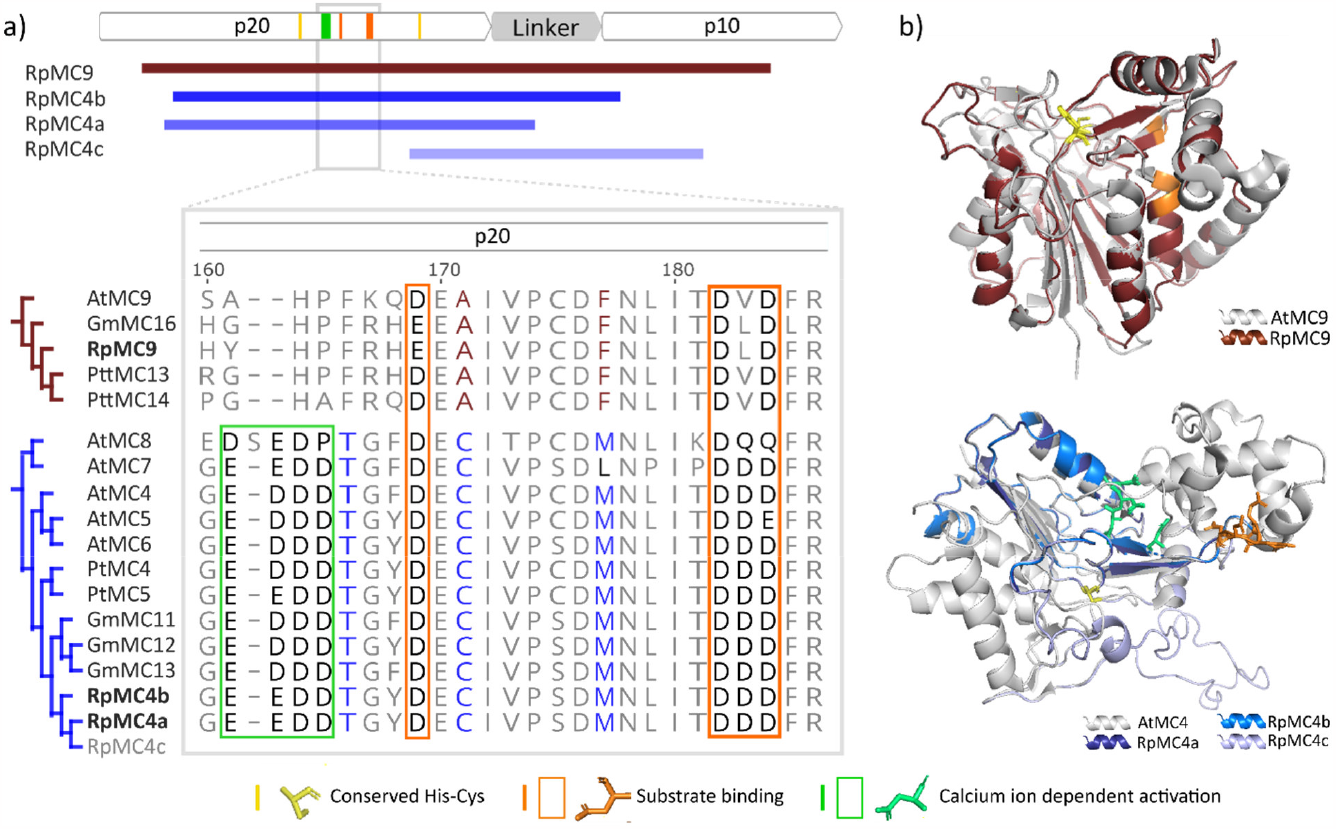
Sequence alignment and homology models of type-II metacaspases (MC-II). a) Protein sequence alignment of MC-II with sequences isolated from *Robinia pseudoacacia (Rp)*. Domain structure elements: white, catalytic domain subunits; grey, linker region. Signature residues of AtMC4 and AtMC9 are colored in blue and red in homologs respectively ^71^. RpMC4a and -b have the negatively charged residues for Ca^2+^-dependent activation as framed in green, but not in RpMC9. b) 3D structural models of RpMC-II based on crystal structures of AtMC4 and AtMC9. Homology models of RpMC-II are superimposed onto the respective templates, showing their overlapping structures. Accessions and quality estimations on homolog models are listed in supplementary materials.

Two partial gene sequences of *VPE* were obtained from *R. pseudoacacia*, designated as *RpVPEβ* and *RpVPEγ* according to their types, and the latter is an identical sequence to an expressed sequence tag (EST; accession BI677606.1) previously reported in Yang et al. (2003; 2004). In addition to proteolytic activities, a list of VPE isoforms exhibits peptide bond ligase ability in neutral pH ^65–68^. Therefore, specific regions in protein sequences identified as the determinants of enzymatic property are compared for functional prediction (Figure 5), including the ligase-activity determinant (LAD) gatekeepers, a plant-specific poly-proline loop (PPL), and marker of ligase activity (MLA) ^65,69,70^. The translated sequences of both *RpVPE* isolated in the present studies contain the signature protease-favored motif in LAD gatekeeper; their MLA are hydrophilic and not truncated, resembling the proteases and bifunctional isoforms.

**Figure 5:**
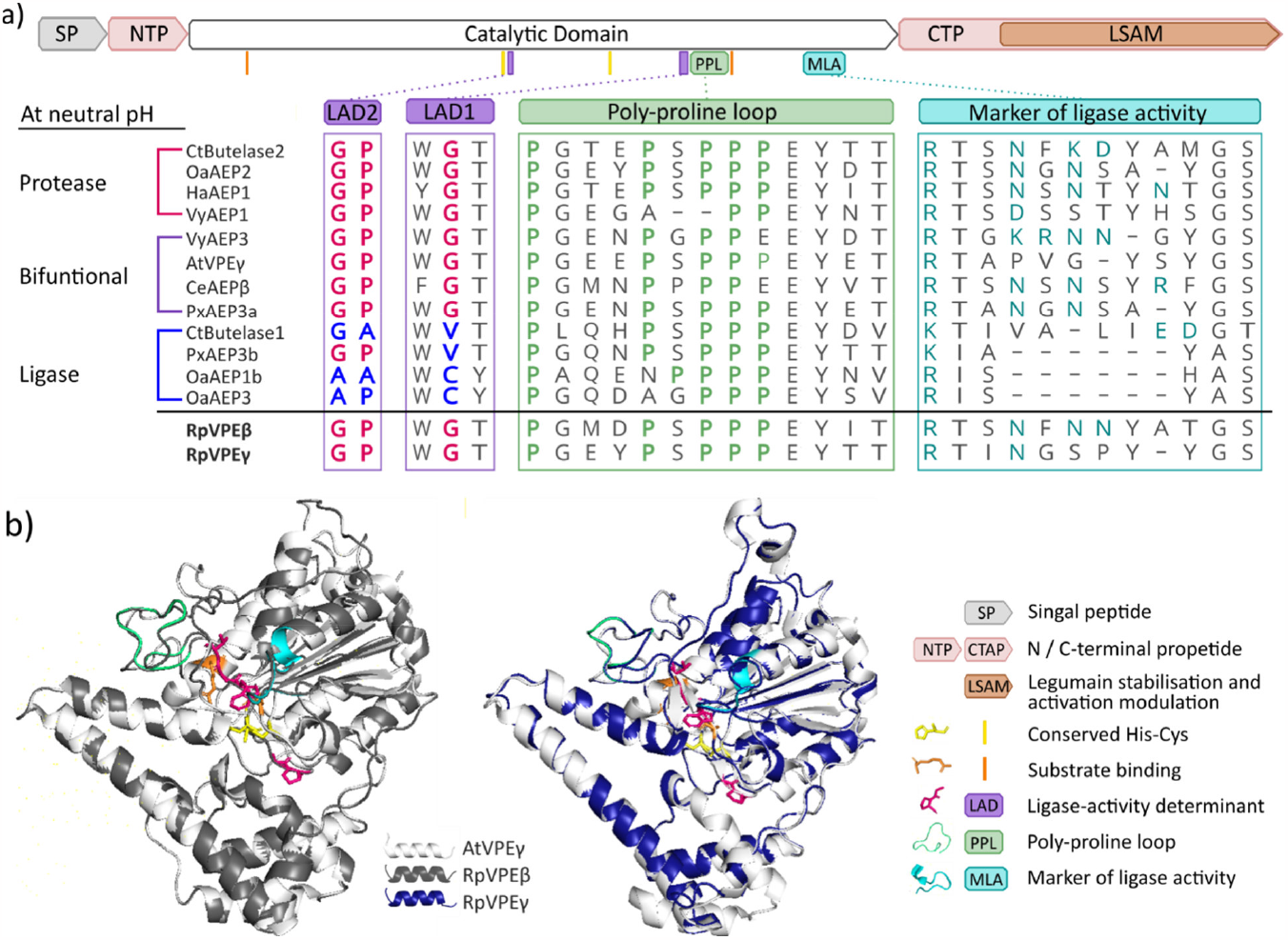
Sequence alignment and homology models of vacuolar processing enzymes (VPE or AEP). a) Sequence alignment of VPEs showing determinants of enzymatic activity with a schematic representation of the domain organization shown on top. Sequences of VPE from *Robinia pseudoacacia* (Rp), *Arabidopsis thaliana* (At), *Canavalia ensiformis* (Ce), *Clitoria ternatea* (Ct), *Helianthus annuus* (Ha), *Oldenlandia affinis* (Oa), *Viola yedoensis* (Vy), *Petunia x hybrid* (Px) are grouped by enzymatic activity at neutral pH. Residues in LAD that favor ligase and protease activities are labeled in magenta and blue, respectively. Prolines in the PPL are labeled green and hydrophilic residues within MLA are in turquoise. RpVPEβ and RpVPEγ are listed in bold under the black line. b) Homology models of RpVPE based on crystal structures of AtVPE as templates. Accessions and quality estimations on homolog models are listed in supplementary materials.

### Gene expression profiles

The indicator of polyphenolic compound production, *RpPAL1*, showed elevated relative expression (RE) in the DX in summer and the TZ during autumn. The flavanols biosynthesis marker gene, *RpCHS3*, showed strong expression exclusively in the TZ between August to November. These results aligned well with previous data derived from a larger number of biological replicates as provided in supplementary materials, where isoforms of the marker genes for polyphenol biosynthesis were assessed.

Throughout the year, fluctuations in RE showed by the *RpVPE* were mild and gradual, while *RpMC-II* presented the contrary in comparison (Figure 6). In the phloem, Ca^2+^-dependent *RpMC-II* showed an upregulated co-expression in July. Gene expression profile in the outer xylem regions is generally in line with extant literature. Following the first sharp increase in both average temperature and sunshine duration of the year, we observed the transcriptional upsurge of *RpMC-II* in the DX in April which gradually declined over the months (Figure 7). *RpMC9* showed a particularly strong elevation, reaching over 200-fold of the reference gene in April and May before a sharp decline. Upregulation of *RpVPEγ* peaked in May, followed by the up-regulation of *RpVPEβ* and *RpPAL*1 in June; *RpPAL1* gradually declined over the summer months. In addition, a transient upregulation of *RpMC9* was seen in July between the MSW to OTZ.

**Figure 6:**
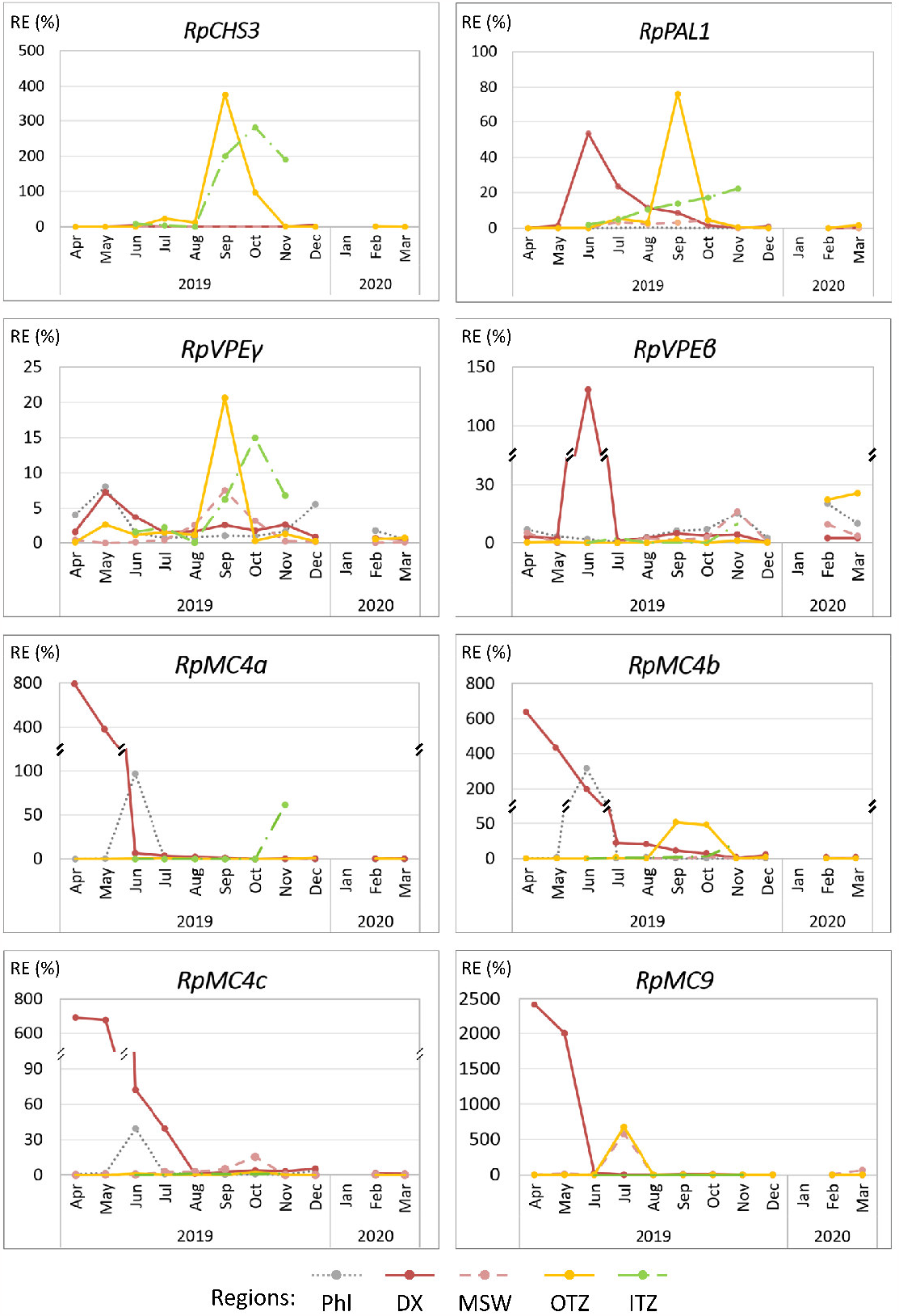
Relative gene expression (RE) of markers for polyphenols biosynthesis and programmed cell death in Robinia stem. Gene expression profiles are relative to the reference gene 18S. Marker genes in *Robinia pseudoacacia (Rp):* Phenylalanine ammonia-lyase 1 (*PAL1*), chalcone synthase 3 (*CHS3*), vacuolar processing enzyme (VPE), and type-II metacaspase (MC-II). Wood Regions: Phloem (Phl), differentiating xylem (DX), middle sapwood (MSW), outer transition zone (OTZ), and inner transition zone (ITZ).

**Figure 7:**
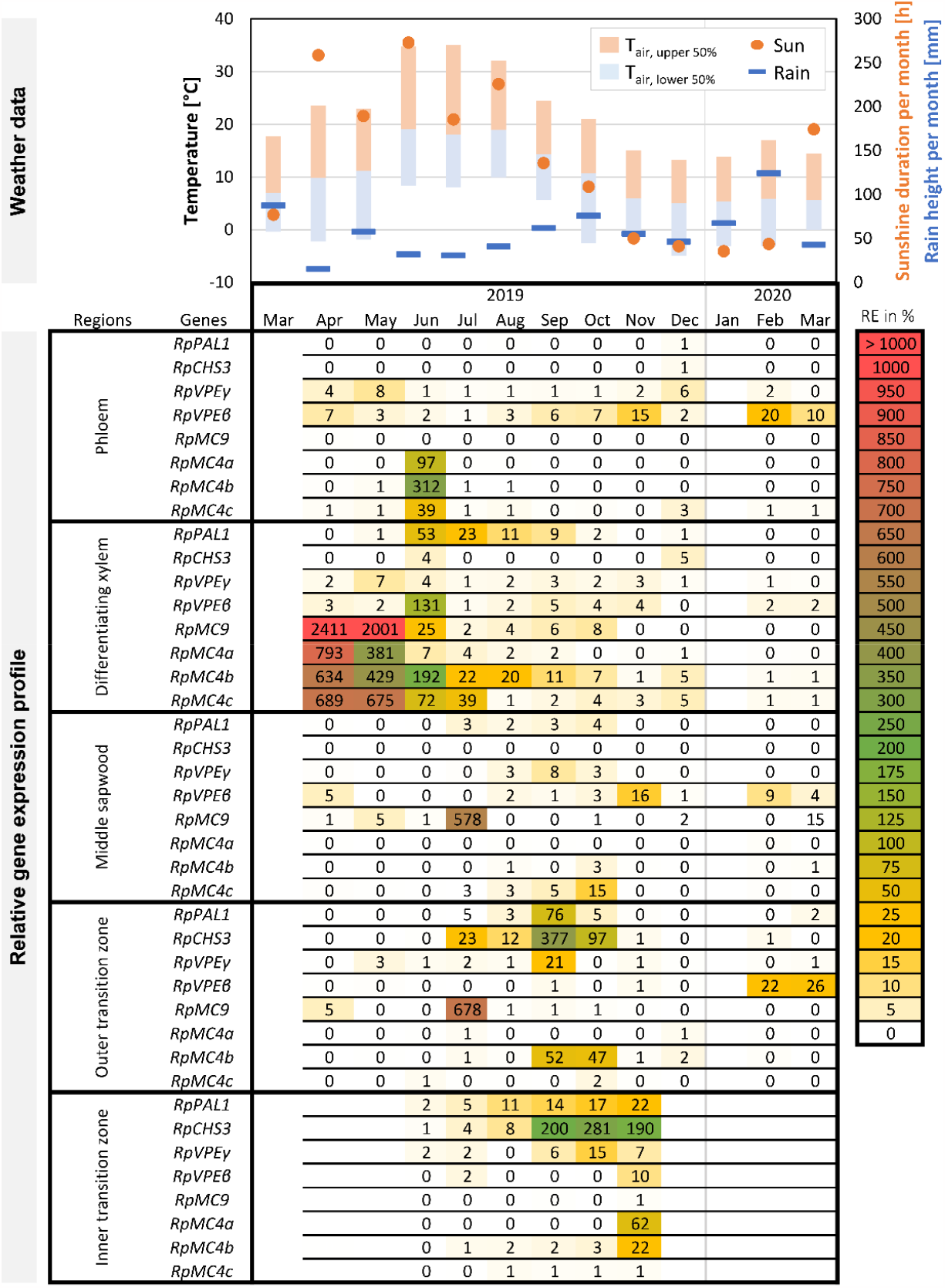
An annual relative gene expression profile of marker genes for polyphenols biosynthesis and programmed cell death in *Robinia pseudoacacia (Rp)* stem, with corresponding weather data. Weather data were sourced from the German Meteorological Service (station 1975 Hamburg-Fuhlsbüttel) with a temporal resolution of 10 minutes. Scale bars indicate air temperature at 2 meters in height; data above and below the mean value are shown in orange and light blue respectively. Gene expression are relative percentage to the reference gene 18S. PAL, phenylalanine ammonia-lyase; CHS, chalcone synthase; VPE, vacuolar processing enzyme; MC, metacaspase.

Results from the transitioning regions present elevated co-expression of the markers for HWF, with similar patterns occurring earlier in the outer region. The synchronized upregulation of *RpPAL1* and *RpCHS3* points towards HWS biosynthesis, which corresponds with the upregulation of the *RpVPEγ* and *RpMC4a* as the co-decline in average temperature and sunshine duration occurred. The peaking RE level of these genes coincided in September, with *RpCHS3* and *RpMC4b* remaining at a high level for another month. At the inner TZ, a similar order of gene expression is shown with a delay of one to two months. November held the highest number of upregulated markers, including *RpVPEβ* and *RpMC4a* that were not presented in the OTZ. Another point to note is that while *MC-IIs* and *RpVPEγ* are synthesized before polyphenols biosynthetic enzymes in the DX, the same is not observed in the TZ.

## Discussions

The current study aimed to identify genetic markers that focus on the PCD aspect during HWF to examine the correlations with HWS biosynthesis and to determine a clear timeline of these occurrences. Our objectives included identifying the reference points for a direct comparison with terminal differentiation observed in other stages of stem development. We isolated members of *MC-II* and *VPE* as PCD markers from *Robinia pseudoacacia* for real-time PCR analysis, and the results demonstrated their suitability as references that correspond with markers for polyphenol biosynthesis. Due to the practical challenges in examining heartwood-forming specimens of unyielding nature, relevant research in the last two decades often makes use of a collection of few samples in spring and autumn. While the current study is bound by similar limitations, results from the reference genes are validated with previous data derived from more biological replicates. Furthermore, our data derived from monthly harvest offers an unprecedented resolution on the fluctuating gene expression throughout the seasonal cycle.

### Vacuole-mediated PCD during heartwood formation

Consistent with previous research, our data demonstrated the upregulation of *RpPAL1* that reflects lignification near the cambium in summer, while expression of *RpCHS3* resonated HWS biosynthesis at the TZ in autumn ^38^. In both cases, corresponding upregulation of *RpVPE* was observed, suggesting their involvement in these processes. γ-type VPE is believed to be the priming hydrolase that promotes tonoplast rupture of the lytic vacuole during developmental PCD, and their higher RE in the DX aligns with previous observations ^56,57^. Our data showed the upregulation of *RpVPEβ* in the DX starting at the same period as *RpPAL1*, suggesting that the involvement of β-type VPE *is* involved during wood formation in addition to *VPEy*. Furthermore, upregulation of *RpVPEβ* is seen in early winter in the MSW, late winter to early spring in the OTZ, and at the late stage of HWF in the ITZ. The upregulation of *VPEγ* in association with HWF has been previously reported in *Taiwania* ^72^ and *Robinia* ^64^, with which our result aligned. Furthermore, our data showed a more prominent upregulation of *RpVPEy* in the TZ than in the DX. Alongside the weather data, it appeared to follow the co-decline in average temperature and sunshine duration in this data set. However, follow-up studies with longitudinal data are necessary to determine a correlating pattern.

### Ca^2+^-dependent MC-II across phloem and xylem

The protein sequences of the isolated RpMC-II contain identical key residues responsible for Ca^2+^-dependent activation and their homolog models predict similar structures, it is therefore safe to assume similar enzymatic properties. In the phloem, we observed the surprising transient up-upregulation of all surveyed Ca^2+^-dependent MC-II, which occurred in the hottest month of the year. This makes an intriguing comparison to the previous report, where only type-I MCs were identified in similar contexts. In *A. thaliana*, AtMC3 had been shown to confer drought tolerance in the phloem by enhancing the differentiation of the metaphloem sieve elements and maintaining higher levels of vascular-mediated transportation ^73^. In the gene expression analysis on the MC family during the season of active xylogenesis in Populus, including both type I and II, distinct upregulation in the phloem was shown with MC-I (*PttMC7, 8, 10*) ^48^; the role of type-II MCs in the phloem await future investigation.

Our data on *RpMC-II* in the DX showing the upsurges of RE in spring is consistent with previous findings. This occurrence followed the first sharp rise in average temperature and sunshine duration of the year. The finding aligns with reports that demonstrated how the timing of cambial reactivation is controlled by temperature, and xylem differentiation starts within 3 or 4 weeks in deciduous trees under natural conditions ^15,74^. It is worth pointing out that while the elevated RE of *RpMC4* in the DX occurred before those of *RpPAL1* and *RpVPE*, it is not the case in the TZ. Instead, their upregulation is shown at the height of RpPAL1’s expression. In addition, there was a positional difference between *RpMC4b* and *RpMC4c*, where the former is observed across the whole transitioning region while *RpMC4c* is only seen in the ITZ.

### Xylem-specific MC9 in aging parenchyma

The lack of *RpMC9* expression in the phloem in our data supports previous findings on its xylem-specificity in the stem ^56,75^. The dramatic upsurge of RE in the DX in spring, along with the Ca^2+^-dependent MC-II, is again in agreement with its involvement in xylem cell differentiation. Diving deeper into the inner wood regions, the transient RE elevation of *RpMC9* between mature sapwood to outer TZ suggests its functions in the aging XP cells. As TEs and fibers complete their terminal differentiation within the first year after cell division at the cambium, the middle sapwood region contains few living cells other than XP cells. However, this remark requires further experiments with detailed separation of cell types using laser microdissection for confirmation. Naturally, the role of MC9 in the different modes of dPCD observed in the various xylem cell types remains an interesting topic for future exploration.

### Timing of heartwood formation and signaling direction

Based on weather data from this harvesting period, the upregulation of markers for xylogenesis followed the first sharp increase in average temperature and sunshine duration of the year, while the markers for HWF corresponded to the co-decline of these conditions. While longitudinal data in follow-up studies are required to confirm a correlated pattern, the observation aligns with previous reports. On the other hand, it is interesting to note that the co-expression of *RpPAL1, RpCHS3, RpVPEγ*, and *RpMC4b* in the outer TZ appeared earlier than that in the inner TZ. One interpretation lies in the tempo-spatial difference previously proposed ^8,10^. A potential explanation is that HWF spans more than one annual cycle, where a majority of XP cells undergo the transformation in the OTZ starting after mid-summer, and the remaining XP cells transform in the year after that begin later in autumn. The idea that different XP cell types are involved also accounts for the differences in the gene expression profile, where *RpMC4a* and *RpVPEβ* were up-regulated in the ITZ but not OTZ. Another interpretation of the observed patterns is a time shift of about a month within the transitioning region. Based on the conclusions drawn from transcriptomic studies on HWF ^9,64,72^, the majority of differentially expressed genes (DEGs) are related to signal transduction, suggesting that the cells located in the innermost part of the trunk are influenced by external environmental conditions. Thus, the expression patterns within the transitioning region in the same year imply that the signals for XP cells to initiate transformation are translocated from the outer region inwards.

## Conclusion

Heartwood Formation stands as a crucial defense strategy in trees, believed to have evolved in response to biotic threats. Its implications for bolstering tree and forest resilience in the face of a changing climate are significant. While the current findings require further validation through studies with larger sample sizes, this research presents an unprecedented preliminary overview of the temporospatial dynamics of HWF, the correlation between the defining cellular processes, and how they correspond to weather conditions. Furthermore, the comprehensive survey of these key cellular processes across different stem regions and throughout the seasons demonstrated that they are shared features in xylem-specific terminal differentiation. This finding establishes genetic markers that can serve as reference points for comparing the molecular regulation underlying stages of stem development, and HWF in various species or under different conditions. Nevertheless, the mechanism governing HWF remains largely enigmatic, influenced by multiple factors. Given that both VPEs and MCs serve as regulators of PCD, their expression profiles offer valuable insights to identify upstream effectors and signaling pathways.

