## Supplementary data for "An annual gene expression profile of markers for programmed cell death and polyphenols biosynthesis across heartwood forming stems in *Robinia pseudoacacia*"

**Content**

1. List of primers for qPCR analysis.
2. Details on the sequences in alignments
3. Homology models: accuracy and quality estimations scores
4. Previous gene expression study on phenylalanine ammonia-lyase and Chalcone synthase in the black locust stem

### 1. List of primers for qPCR analysis.

| Gene name | qPCR Primers | | | | Amplicon | |
|---|---|---|---|---|---|---|
| | Accession | Sequence (5' to 3') | Ta (°C) | Eff. | Size (bp) | Tm (°C) |
| *Rp18S* | EF494737.1 | F: GCTTTTAGGACTCCGCTGG<br>R: GGTAAGTTTCCCCGTGTTGA | 60 | 2.02 | 155 | 83.3 |
| *RpPAL1* | EU650627 | F: GGAATGCCTTGGGGAGTG<br>R: CGACGCTGTATTGATTTGCC | 60 | 2.04 | 147 | 78.6 |
| *RpCHS3* | EU650632 | F: ACCACAGGTGAAGGACTCG<br>R: GATAAAATCCAGGTCCAAAGC | 60 | 2.03 | 127 | 80.6 |
| *RpMC9* | OR640705 | F: AACATTGAGCTCCTCACTGA<br>R: TGGCGGAAAGGATGGTAATG | 62 | 1.96 | 182 | 82.2 |
| *RpMC4a* | OR640708 | F: CAATACCCCTAGAGGTGTTAGGATC<br>R: CATCATCCCTGCGTCCATGA | 64 | 1.94 | 236 | 82.1 |
| *RpMC4b* | OR640707 | F: AGATGTGCTGTTCGTCCATTACA<br>R: GACATCGTCTACAAATTCCCTGAAAT | 65 | 1.91 | 151 | 80.8 |
| *RpMC4c* | OR640706 | F: GAAAATGTTAAGGAGCAGATAGGAGAG<br>R: CATCATCCCTGCGTCCATGA | 64 | 2.07 | 175 | 80.8 |
| *RpVPEβ* | OR640709 | F: ATCCTGCCACCGTGAACTTT<br>R: CTGTGCTTCACTGTCTCCGT | 60 | 1.96 | 172 | 80.7 |
| *RpVPEγ* | OR640710 | F: AAGCAGTCAACCAACGGGAT<br>R: GTTGAGCACTTCTGGACCCT | 60 | 1.94 | 197 | 80.1 |

F = forward; R = reverse; Ta = annealing temperature; Eff. = Efficiency

### 1. Details on the sequences in alignments

Sequences in Figure 1. Protein sequence alignment of metacaspases (MC).

| | Species | Accession |
|---|---|---|
| AtMC9 | *Arabidopsis thaliana* | Q9FYE1 |
| GmMC16 | *Glycine max* | I1N8J0 |
| RpMC9 | *Robinia pseudoacacia* | Translated OR640705 |
| PttMC13 | *Populus tremula x tremuloides* | Potri.016G024500.1 |
| PttMC14 | *Populus tremula x tremuloides* | Potri.006G026500.1 |
| AtMC8 | *Arabidopsis thaliana* | Q9SA41 |
| AtMC7 | *Arabidopsis thaliana* | Q6XPT5 |
| AtMC4 | *Arabidopsis thaliana* | O64517 |
| AtMC5 | *Arabidopsis thaliana* | O64518 |
| AtMC6 | *Arabidopsis thaliana* | O64519 |
| PtMC4 | *Populus trichocarpa* | XP_002316158.1 |
| PtMC5 | *Populus trichocarpa* | XP_002311278.2 |
| GmMC11 | *Glycine max* | I1KW32 |
| GmMC12 | *Glycine max* | I1MIC1 |
| GmMC13 | *Glycine max* | I1KW34 |
| RpMC4b | *Robinia pseudoacacia* | Translated OR640707 |
| RpMC4a | *Robinia pseudoacacia* | Translated OR640708 |
| RpMC4c | *Robinia pseudoacacia* | Translated OR640706 |

Sequences in Figure 2. Sequence alignment of vacuolar processing enzymes (VPE or AEP).

| | Species | Accession |
|---|---|---|
| CtButelase2 | *Clitoria ternatea* | KR912009 |
| OaAEP2 | *Oldenlandia affinis* | KR259378 |
| HaAEP1 | *Helianthus annuus* | AIZ09514 |
| VyAEP1 | *Viola yedoensis* | TR78040 |
| VyAEP3 | *Viola yedoensis* | TR35538 |
| AtVPE | *Arabidopsis thaliana* | Q39119 |
| CeAEP | *Canavalia ensiformis* | P49046 |
| PxAEP3a | *Petunia x hybrid* | MG720072 |
| CtButelase1 | *Clitoria ternatea* | KF918345 |
| PxAEP3b | *Petunia x hybrid* | MG720076 |
| OaAEP1b | *Oldenlandia affinis* | KR259377 |
| OaAEP3 | *Oldenlandia affinis* | KR259379 |
| RpVPEγ | *Robinia pseudoacacia* | Translated OR640710 |
| RpVPEβ | *Robinia pseudoacacia* | Translated OR640709 |

### 2. Homology models: accuracy and quality estimations scores

**RpMC4a**

GMQE: 0.

GMEANDisCo Global: 0.78 ± 0.05

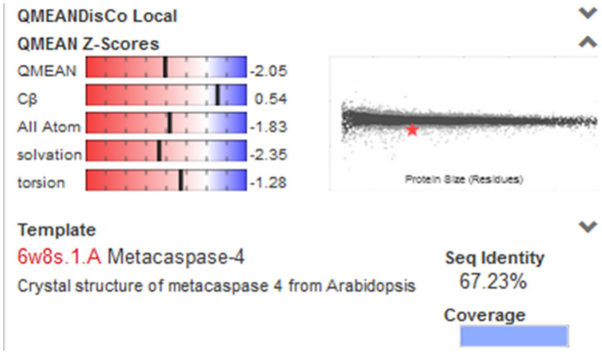


**RpMC4b**

GMQE: 0.69

GMEANDisCo Global: 0.79 ± 0.08

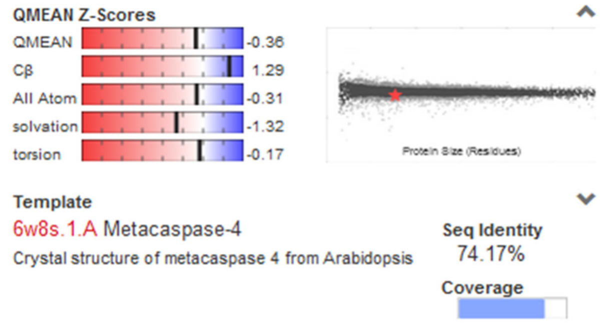


**RpMC4c**

GMQE: 0.36

GMEANDisCo Global: 0.46 ± 0.08

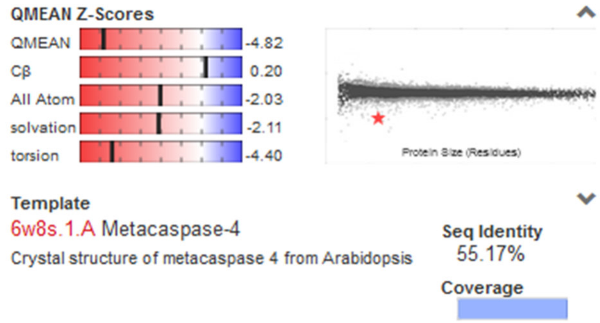


**RpMC9**

GMQE: 0.81

GMEANDisCo Global: 0.78 ± 0.05

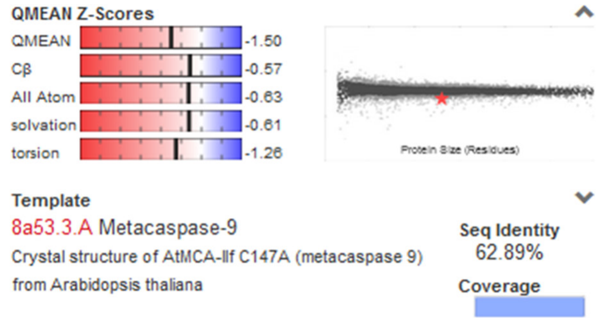


**RpVPEγ**

GMQE: 0.88

GMEANDisCo Global: 0.81 ± 0.05

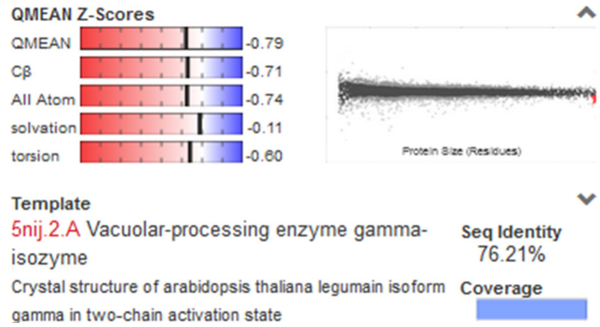


**RpVPEβ**

GMQE: 0.87

GMEANDisCo Global: 0.83 ± 0.05

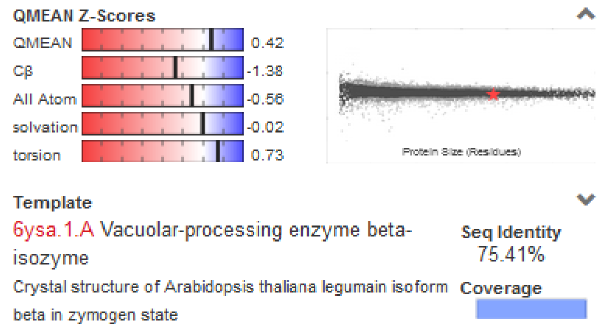

### 3. Previous gene expression study on phenylalanine ammonia-lyase and Chalcone synthase in the black locust stem

#### Materials and methods

Differences between the current experiment in the main article and previous data produced in the same laboratory and with the same equipment:

<table>
<tr><th></th><th>Previous data [1]</th><th>Current experiment</th></tr>
<tr><td>Plant materials</td><td>Specimen<br>Stem wood from the main struck<br>(Tree ages: 8 to 20 years old; trunk diameter at breast height: 10 to 20 cm)<br><br>Tree location<br>Forest sites in Hamburg-Bergedorf and Reinbek, Germany<br><br>Harvest time and amount<br>3 trees in each harvest<br>Spring: Early May, 2006<br>Midsummer: Early Jul, 2006<br>Late summer: End of Aug, 2006<br>Autumn: End of Oct, 2006<br>Late Autumn: End of Nov, 2006</td><td>Specimen<br>Stem wood from the branches<br>(Tree ages: about 25 years old; trunk diameter at breast height: 20 to 40 cm)<br><br>Tree location<br>Arboretum of the Thünen Institute in Hamburg-Bergedorf, Germany<br><br>Harvest time and amount<br>1 branch in each harvest<br><br>First Fridays in the months between April 2019 to March 2020</td></tr>
<tr><td>RNA isolation</td><td>Reagent<br>RNeasy Plant Mini Kits, Qiagen</td><td>Reagent<br>RNeasy Plus Universal Kits, Qiagen</td></tr>
<tr><td>DNase treatment</td><td>Reagent DNase-Verdau Deoxyribonuclease I (DNase I), Fermentas</td><td rowspan="2">Combined step<br><br>Reagent<br>QuantiNova Reverse Transcription Kit, Qiagen</td></tr>
<tr><td>Reverse transcription</td><td>Reagent<br>SuperScript® III first strand synthesis kit, Invitrogen</td></tr>
<tr><td>qPCR</td><td>Reagent<br>Brilliant® II SYBR® Green QPCR Master Mix, Stratagene (Agilent Technologies)</td><td>Reagent<br>Quantinova qPCR SYBR Green Master Mix (QIAGEN)</td></tr>
<tr><td colspan="3">1. Lange, H. Molekulare Grundlagen von Verfärbungsprozessen im Holz der Robinie (Robinia pseudoacacia L.): Genexpressionsstudien an Schlüsselgenen der Flavonoidsynthese. (Universität Hamburg, 2009).</td></tr>
</table>

### Results

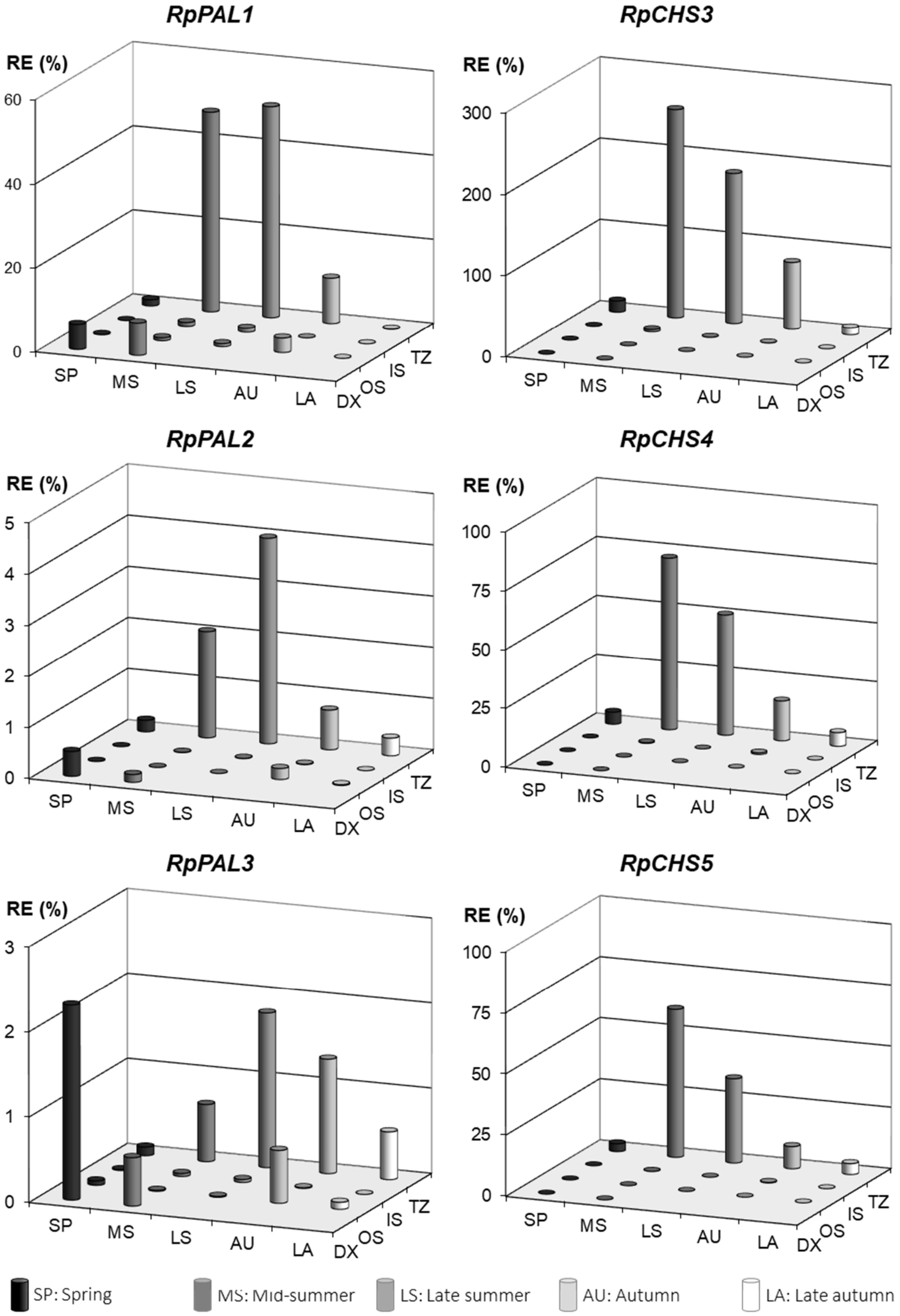


Figure S1: Relative gene expression (RE) of phenylalanine ammonia-lyase (PAL) and chalcone synthase (CHS) isoforms in healthy black locust stem subdivided by seasons and regions. RE values are relative to the reference gene *Rp18S*. Wood Regions: Differentiating xylem (DX), outer sapwood (OS), inner sapwood (IS), and transition zone (TZ).
